# Catalytic degradation of circulating targets with FcRn-mediated cycling LYTACs

**DOI:** 10.1101/2025.01.12.632472

**Authors:** Christos Kougentakis, Matthew J. Shurtleff, Rachel M. Lieser, Tao Chen, Christina L. Collins, Isabel E. Johnson, Siyi Gu, Keith T. Akama, Kyle H. Doran, Cyra K. Fung, Darin J. Hildebrandt, Rozhan Khaleghi, Diana Li, Steve Rank, Thomas-Toan Tran, Richard Yau, Ryan W. Holly, Christopher Langsdorf, Darrin A. Lindhout, Sarah M. McWhirter, Jeffrey S. Iwig, Jason G. Lewis, Steven T. Staben, Eric D. Turtle, Nicholas A. Lind

**Author notes:** These authors contributed equally: Christos Kougentakis, Matthew J. Shurtleff.

## Abstract

Circulating proteins are common targets for the discovery of occupancy-based inhibitors including monoclonal antibodies. Effective inhibition of target pathogenicity with blocking approaches, however, is often challenged by target parameters that lead to insufficient occupancy and/or incomplete pharmacology limited by only single site binding. Extracellular targeted protein degradation approaches, such as lysosomal targeting chimeras (LYTACs), offer an opportunity to minimize these challenges by an event-driven mechanism that selectively, thoroughly and irreversibly eliminates drivers of disease. First generation LYTACs, designed to traffic to the lysosome, show limited durability since the therapeutic is degraded along with the target protein of interest. Here we describe cataLYTACs, which overcome this limitation by combining stabilized asialoglycoprotein (ASGPR) ligands, pH-sensitive target binding and recycling via the neonatal Fc receptor (FcRn). These cataLYTACs degraded superstoichiometric levels of a target protein, IgE, in vitro and demonstrated deep and sustained clearance of human IgE in mouse models. In non-human primates, cataLYTACs resulted in >98% clearance of circulating endogenous IgE for 2 weeks and outperformed the standard of care blocking antibody, omalizumab (Xolair®), in both free IgE elimination and duration of action. CataLYTACs represent a new therapeutic modality for a wide range of disease states driven by circulating factors, with the potential for superior efficacy and duration of action compared to traditional inhibitors.

## Introduction

The serum proteome is a complex mixture of >2000 proteins originating from all tissues with circulating concentrations ranging over ∼10 orders of magnitude and protein half-lives ranging from minutes to weeks^1,2^. This diversity is maintained, in part, through processes that can selectively shorten (*e.g.*, proteases, scavenger receptors) or lengthen (*e.g.,* recycling pathways, extracellular chaperones) the half-life of individual proteins. Secreted circulating proteins are critical mediators of health and disease but high abundance targets (*e.g.,* apolipoproteins, complement proteins, immunoglobulins, clotting factors) pose a challenge to achieve therapeutic inhibition using conventional occupancy-based blocking approaches (*e.g.*, antibodies, small molecules).

Extracellular targeted protein degradation (eTPD) is an emerging therapeutic field employing various modalities to achieve the targeted degradation of plasma membrane and secreted soluble proteins to treat disease^3,4^. Targeted degradation, as opposed to inhibition, operates through a differentiated principle of event-driven pharmacology in which a binding event triggers active elimination of the target^5-7^. These approaches employ heterobifunctional molecules to induce proximity between a target protein and a cell surface protein that directs the target for degradation in the lysosome. Lysosome targeting chimeras (LYTACs) and other molecular degraders of extracellular proteins co-opting the asialoglycoprotein receptor (ASGPR) have been described that mediate rapid clearance of circulating target proteins in the liver by ASGPR-expressing hepatocytes in <6 hrs^8-10^. On mechanism ASGPR-mediated clearance for these modalities, however, presents challenges with respect to both efficiency and durability. The therapeutic is degraded with similar rapidity in the presence or absence of the target protein of interest. This characteristic necessitates a large excess of therapeutic and substantially minimizes the ability to degrade regenerated target without high dosing frequency.

Given a lack of need for constant target occupancy, an extracellular targeted protein degradation (eTPD) approach is particularly attractive for high abundance circulating pathogenic target proteins that are not amenable to inhibition with monoclonal antibodies due to dosing limitation. For example, omalizumab (Xolair^®^) is commonly prescribed for treatment of IgE-mediated allergic diseases, however, dosing limitations restrict its use as a monotherapy in patients with high circulating IgE levels, such as severe allergic asthma and other atopic indications^11,12^. Unlike inhibitors, which require constant occupancy, an eTPD approach could theoretically enable removal of a superstoichiometric molar concentration of target to drug via a catalytic degradation mechanism in which the drug facilitates target degradation without being consumed in the process. This mechanism allows each drug molecule to degrade multiple target molecules. Proteolysis targeting chimeras (PROTACs) have been shown to exhibit catalytic degradation properties via the proteasome^5^. However, circulating soluble proteins are generally not amenable to a PROTAC approach since they are co-translationally translocated to the endoplasmic reticulum for secretion and therefore not accessible to the ubiquitin-proteasome degradation pathway in the cytoplasm.

Here we describe LYTACs that combine rapid ASGPR-dependent uptake with pH-sensitive target binding and recycling to achieve catalytic degradation of IgE. These cataLYTACs^TM^ demonstrate degradation of superstoichiometric levels of IgE in vitro as well as deep and sustained elimination of circulating IgE in mouse and non-human primate models.

## Results

### Deep and rapid depletion of target molecules with O-linked triGalNAc LYTACs

The use of 1-O-glycosidic linked GalNAc (N-acetylgalactosamine) derived groups to direct binding to ASGPR and subsequent endocytosis into the liver has been demonstrated for both drug delivery and degradation modalities^8-10,13^. Previously described monomeric GalNAc ligands for ASGPR are low affinity (K_D_∼40 µM)^14^ and require multimeric display to achieve high affinity through avidity and preferred geometric presentation to the trimeric ASGPR complex^15,16^. A schematic representation of IgE-targeting LYTACs utilized in this study is displayed in Figure 1a. Human IgG1 monoclonal antibodies that bind to IgE were expressed with an introduced cysteine (L443C) to allow site specific conjugation of ASGPR targeting ligands. The small molecules consisted of a core linker for the mono-, bi-, and tri-antennary display of ASGPR binding groups (Fig 1a - Ligands), and a spacer and cysteine reactive moiety for attachment to an antibody (Fig 1a - LYTACs). Ligand (**Mono 1**, **Bi 1** and **Tri 1**) binding to ASGPR was assessed using a fluorescence polarization-based competitive binding assay. The expected improved ASGPR binding affinity with increasing ligand valency was observed (Extended Data Fig 1a-b).

**Fig. 1.**
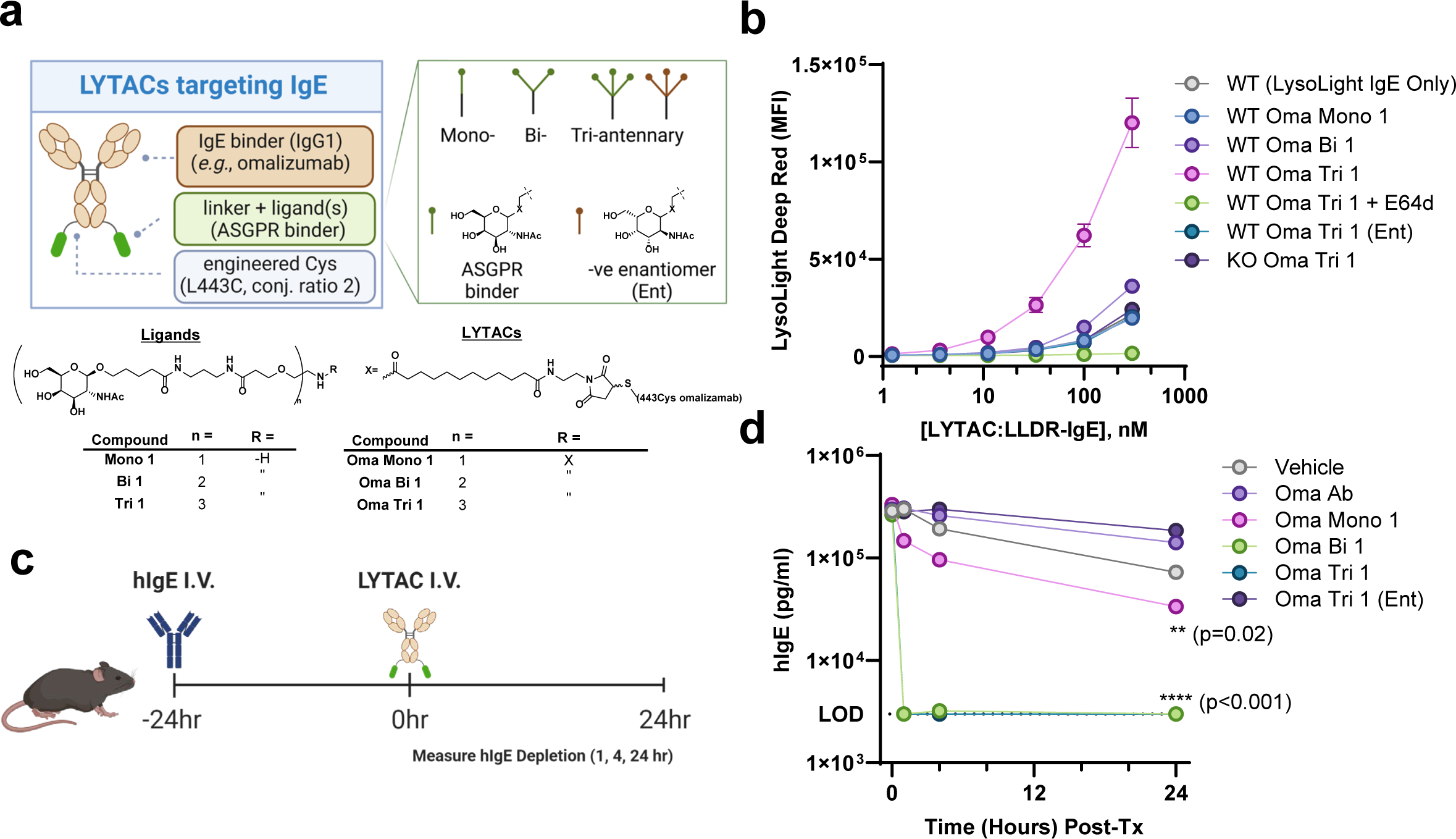
ASGPR targeted LYTAC molecules allow for rapid and deep depletion of circulating targets. **(a)** Schematic showing components of ASGPR-targeted a-IgE LYTAC molecules; structure of ligands and LYTAC compounds. **(b)** Lysosomal delivery and catabolism of LysoLight Deep Red labelled-IgE molecules in LYTAC treated HepG2 WT and ASGPR1/2 KO cells. Enantiomer of trivalent GalNAc labelled (Ent). Cathepsin protease inhibitor E64d was added where indicated. **(c)** Schematic of *in vivo* pharmodynamics study to determine LYTAC depletion efficiency. C57BL/6 mice were dosed with 1 mg/kg human IgE intravenously 24 hours before being dosed with 5 mg/kg test articles subcutaneously. **(d)** *In vivo* clearance of human IgE by LYTAC molecules (n=3/group). Statistical analysis: 2way ANOVA with

Next, omalizumab-based LYTACs incorporating the aforementioned ligands (**Oma Mono 1**, **Oma Bi 1** and **Oma Tri 1**) were evaluated in an assay measuring IgE uptake and lysosomal degradation. ASGPR-expressing HepG2 cells were incubated with LYTAC and LysoLight^TM^ Deep Red-conjugated IgE (IgE-LLDR) and fluorescence was measured by flow cytometry (Fig 1b). LysoLight Deep Red is a probe that fluoresces only upon cleavage by lysosome resident cathepsin proteases to provide a surrogate measure for target degradation^17^. **Oma Tri 1** and **Oma Bi 1** LYTACs promoted increased uptake/degradation over the IgE-only control. Activity of the **Oma Mono 1** was indistinguishable from a negative control conjugate (**Oma Tri Ent.**), where the binding elements were replaced with enantiomeric GalNAc groups incapable of binding to ASGPR (Fig 1b). LYTAC treatment in ASGPR1/2 knockout cells did not increase catabolism over the IgE-LLDR only control and no signal was observed in the presence of the cathepsin protease inhibitor E64d (Fig 1b).

These results demonstrate LYTAC and ASGPR-dependent lysosomal degradation of IgE-LLDR in vitro. In vivo activity of these LYTACs was determined in a mouse pharmacodynamic model with exogenous administration of hIgE followed by intravenous treatment with LYTAC (Fig 1c). **Oma Bi 1** and **Oma Tri 1** LYTACs depleted over 90% of the hIgE by one hour, while the **Oma Mono 1** cleared the target more slowly and with a lower depth of depletion (∼50% in 24 hours) (Fig 1d). The ASGPR non-binding control (**Oma Tri Ent.**) matched the activity of the parent antibody alone. Taken together, LYTACs bearing O-linked ligands promote ASGPR-dependent uptake and degradation of IgE in vitro and rapid clearance of exogenously administered IgE in vivo. We next sought to improve the durability of LYTACs by eliminating predicted catabolic liabilities within the ligands.

### Engineered ASGPR ligands with increased affinity and stability for enhanced target degradation and increased duration of action

CataLYTACs are envisioned to undergo multiple internalizations to an endocytic compartment with potentially long residence times in both extracellular and endocytic environments. Acid hydrolases, such as cathepsins and gylcosidases, within the endolysosomal pathway mediate catabolism of endocytosed glycoproteins^18,19^. We therefore hypothesized that removing bonds constituting potential substrates for endolysosomal enzymes within the exposed Tri linker-ligand (**Oma Tri 1)** would result in stabilized LYTACs capable of retaining activity through multiple rounds of cycling (Fig 2a, potential metabolic soft spots highlighted in red).

**Fig. 2.**
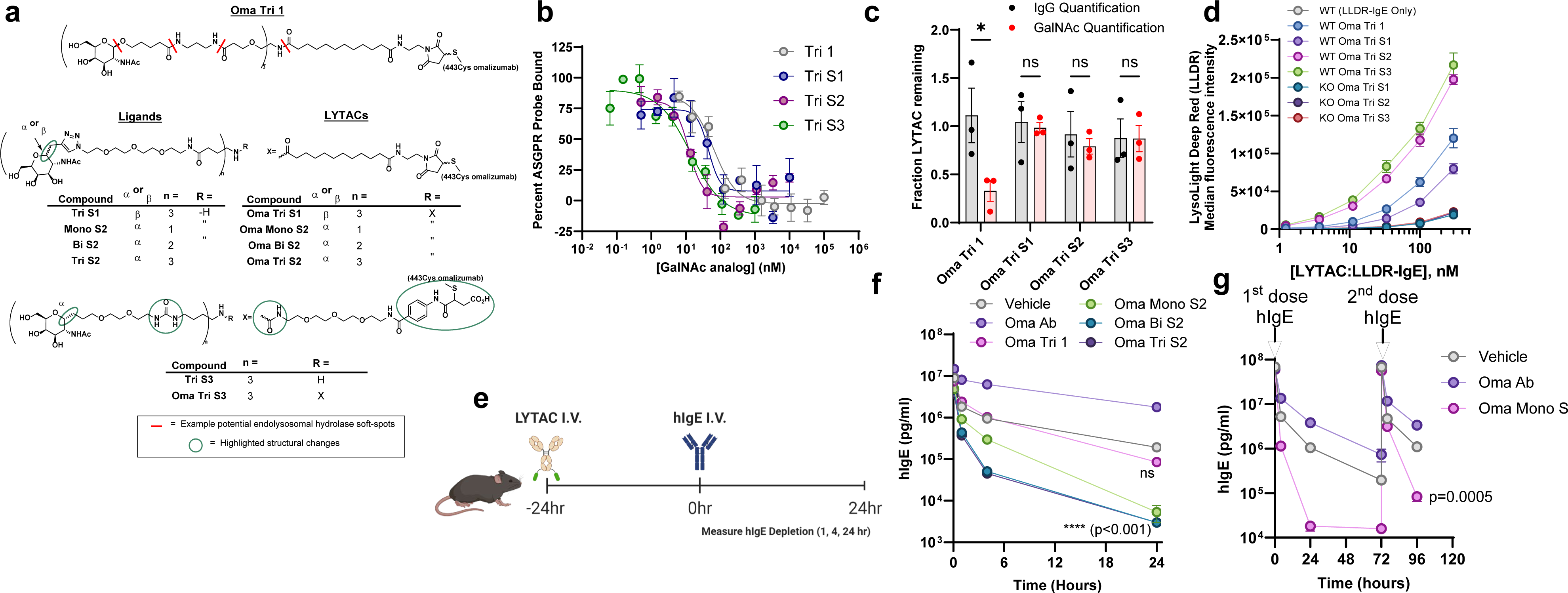
Optimized ASGPR ligands improve LYTAC activity and stability. **(a)** Chemical structures of ASGPR-binding linker-ligands. Potential catabolic sites in Compounds 1 are labelled in red hashes **(b)** Fluorescence polarization competition binding assay for ligands binding to ASGR1. **(c)** Tritosome stability assay of Oma LYTACs with different ASGPR ligands. LYTACs were incubated with tritosomes for 0 or 3 hours, and concentrations were measured using a-IgG or ASGPR ELISAs. Fraction of LYTAC remaining after 3 hours, quantifying both IgG and GalNAc concentrations are shown. **(d)** IgE degradation assay with LysoLight Deep Red labelled IgE in HepG2 WT and ASGPR1/2 KO cells. **(e)** Schematic summarizing dosing schedule for in vivo delayed dosing mouse studies. **(f)** Delayed challenge experiment showing depletion of 1 mg/kg intravenously administered hIgE by stabilized LYTAC molecules (5 mg/kg, subcutaneous, for all experiments)(n3/group). Statistical analysis: 2way ANOVA, significance calculated compared to Vehicle group. **(g)** Delayed challenge experiment as in (e), but with a second 5mg/kg dose of hIgE intravenously administered 72 hours after an initial 5mg/kg hIgE dose (n=2-3/group). Statistical analysis: Unpaired t test at 96hr timepoint, significance calculated compared to Vehicle group.

We prepared a series of stabilized derivatives to eliminate potential catabolic sites in the exposed chemical ligand. Key modifications to **Oma Tri 1** are highlighted in Fig. 2a and chemical structures are presented in Extended Data Fig 5. We first replaced the 1-O-glycosidic linkage used above with 1-carbon substitution (C-glycoside derivatives S1, S2; β and α stereochemistry, respectively). Trivalent ligands incorporating 1-C-α configuration (**Tri S2**) were observed to have superior affinity compared to the 1-C-β configuration (**Tri S1**, Fig 2b). 1-C-α linked ligands were also examined in a mono, bi, and tri-antennary display (Extended Data Fig 1a-c, **Mono S2**, **Bi S2**, **Tri S2**) and these analogs followed affinity trends observed for other GalNAc-based ASGPR binders: Tri>Bi>>Mono. Further modification (**Tri S3**) replaced potentially labile amides in the linker and conjugation through an N-aryl-maleimide capable of undergoing mild, one pot addition and ring-opening to ensure stability at the site of conjugation^20^. **Tri S3** demonstrated comparable binding affinity with **Tri S2** to ASGPR (Fig 2b).

We next evaluated stability associated with the above structural modifications by incubating the corresponding LYTACs with tritosomes to simulate endolysosomal compartment activity^21,22^. **Oma Tri 1**, **Oma Tri S1**, **Oma Tri S2** or **Oma Tri S3** were incubated with rat liver tritosomes for 3h and total IgG and ASGPR ligand levels were measured by ELISA (Fig 2c). LYTACs with stabilized ligands (**Oma Tri S1, S2 and S3**) were resistant to catabolism by hydrolases present in the tritosome fraction compared to the parent unstabilized **Oma Tri 1** (Fig 2c). Furthermore, the **Oma Tri S2** and **Oma Tri S3** LYTACs demonstrated improved in vitro activity relative to **Oma Tri S1** or **Oma Tri 1** (Fig 2d).

To determine whether the improvements in ASGPR ligand stability and affinity translated to improved duration of action in vivo, we developed a pharmacodynamic protocol that requires LYTACs to remain active for a defined period of time before introduction of the soluble target (Figure 2e). **Oma Tri 1,** which demonstrated robust activity when administered after the soluble target (Figure 1e), showed expected minimal activity when dosed 24 hours prior to target in this “delayed challenge” format. (Figure 2f). By contrast, **Oma Mono-**, **Bi-** or **Tri S2** LYTACs, regardless of valency, resulted in rapid clearance of hIgE under the same conditions (Figure 2f). These initial results motivated us to further explore the duration of action for stabilized LYTACs by increasing the delay between LYTAC and target administration up to 7 days, and by challenging with repeated rounds of soluble target (Figure 2g, Extended Data Fig 1d). The stabilized LYTACs were capable of target clearance in each of these scenarios, demonstrating enhanced duration of action and thus potential for catalytic activity.

### LYTACs with next generation engineered ASGPR ligands are capable of cycling activity through multiple cellular pathways

The improved durability observed with **Oma Tri S2** LYTAC in vivo could be explained by the ability to survive multiple rounds of internalization and recycling. Accordingly, in vitro cycling assays were developed to directly monitor LYTAC recycling (Fig 3a). Cells were incubated with Alexa Fluor488 (A488) labeled LYTACs (uptake), transferred to ice to temporarily halt trafficking, stringently washed to remove surface bound LYTAC and then incubated with fresh media containing Alexa Fluor647 (A647) labeled detection reagent (anti-human IgG antibody in Fig 3b, human IgE in Fig 3c-f) to detect any LYTAC molecules recycling to the cell surface. We first evaluated the role of ASGPR and FcRn in LYTAC recycling using a time course assay in which the recycling signal (Alexa 647) was normalized to the uptake signal (Alexa 488) to obtain a relative measure of cycling efficiency. The addition of excess high affinity competitor ASGPR ligand (**Tri S3**) during the cycling period of the assay reduced, but did not eliminate, the efficiency of **Oma Tri S2** cycling (Fig 3b). We hypothesized that FcRn, a well-established mediator of Ab recycling expressed in HepG2 cells and hepatocytes, was responsible for the remaining cycling activity^23,24^. In FcRn knockout HepG2 (FcRn KO) cells, cycling was reduced relative to wild type cells, however LYTAC accumulation in the cells was increased consistent with elimination of a recycling pathway (Fig 3b, Extended Data Fig 2a-b). Residual recycling in the FcRn KO line was abolished in the presence of competitive ASGPR ligand (**Tri S3**), demonstrating that FcRn and ASGPR contribute to LYTAC cycling in vitro (Fig 3b).

**Fig. 3.**
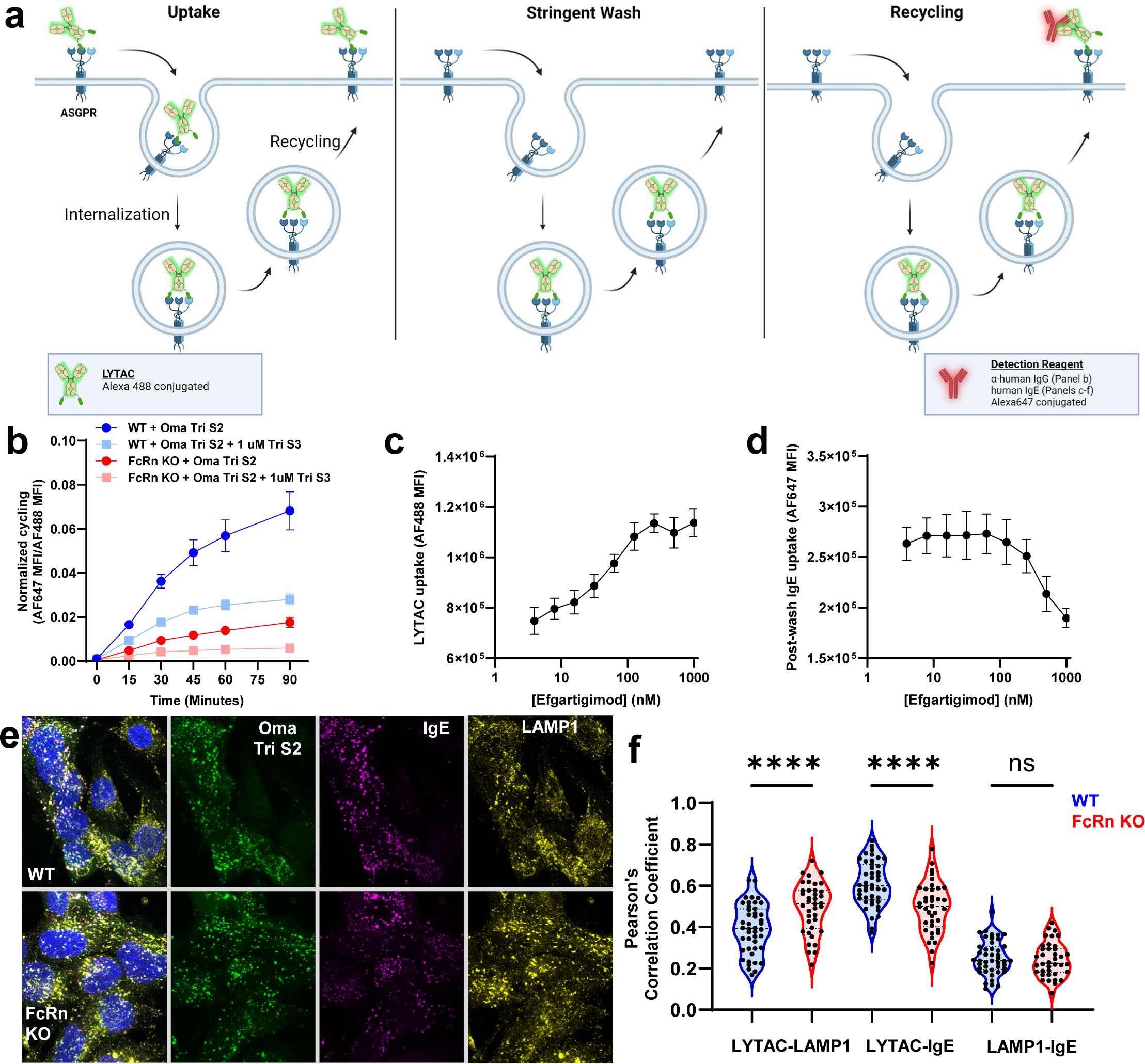
FcRn is necessary for efficient cycling of LYTAC molecules. **(a)** Schematic representation of recycling assay. **(b)** Recycling assay measuring Oma Tri S2 in HepG2 WT and FcRn KO cells, with or without a high affinity competitor ASGPR ligand (Tri S3) introduced during the cycling period. LYTAC recycling signal (Alexa Fluor 647 labelled α-IgG) normalized to LYTAC uptake signal (Alexa Fluor 488 conjugated to LYTACs). **(c-d)** LYTAC uptake (c) and cycling (d), measured by target Alexa Fluor 647 labelled IgE uptake after 4 hours in the presence of the FcRn inhibitor efgartigimod. **(e-f)** Immunofluorescence of Alexa Fluor 488-labelled cycling LYTACs (green) and Alexa Fluor 647-labelled IgE (magenta) relative to LAMP1-positive vesicles (yellow) in HepG2 cells. Quantification of colocalization shown in (f). Tukey’s multiple comparisons test, significance calculated compared to Oma Ab group at 24hr.

To confirm the role of FcRn in LYTAC recycling, the FcRn inhibitor, efgartigimod, was added during the uptake phase of the experiment. Efgartigimod treatment resulted in dose-dependent accumulation of intracellular LYTAC (Fig 3c) and reduced recycling (Fig 3d). This finding was recapitulated with genetic knock out of FcRn (Extended Data Figure 2c-e). Utilizing human IgE as the detection reagent ensured that recycled LYTAC can still bind target. Finally, the recycling assay was repeated using immunofluorescent confocal microscopy as the primary readout to observe localization of the internalized LYTAC molecule relative to both the IgE internalized after cycling and a lysosomal marker (LAMP1). The LYTAC showed increased colocalization with lysosomes and decreased colocalization with IgE in FcRn KO cells compared to WT (Fig. 3e-f). These results show that although FcRn KO cells accumulate more LYTAC, a large proportion is trafficked to the lysosome and consequently unable to recycle and bind extracellular target. Consistent with this interpretation, FcRn inhibition increased LYTAC-LLDR signal in HepG2 cells cotreated with **Oma Tri S2**-LLDR and efgartigimod, confirming that FcRn inhibition increased LYTAC delivery to the lysosome (Extended Data Fig 2f). Taken together, these results demonstrate that FcRn plays a critical role in rescuing cycling LYTACs from lysosomal degradation and is a likely driver of extended durability observed with stabilized ligands in vivo.

### Catalytic degradation of IgE in vitro with pH-sensitive cycling LYTACs

Although the next generation ASGPR ligands allow for efficient LYTAC cycling, we observed relatively low colocalization of IgE with lysosomal markers following post-cycling internalization (Fig 3f). This observation suggested that IgE is cycling with the LYTAC, resulting in inefficient delivery to the lysosome. To improve lysosomal delivery of the target, we engineered pH-sensitive IgE release at endosomal pH (∼ 6.0). Using a limited histidine mutagenesis screen of candidate residues in the CDR of omalizumab (Oma) close to the IgE binding interface, we found substitutions that increased pH-sensitive binding (Extended Data Fig 3a-b). When combining two of these substitutions (S35H-LC, Y57H-LC; Oma pH-S), we observed pH-sensitive binding with increased target release at pH 6.0 (Fig. 4a, Extended Data Fig 3c, e). To test the generality of the benefits of a pH-sensitive binder for a given target to cycling LYTACs, we also tested another IgE binder, ligelizumab (Lige)^25^, as well as a previously described pH-sensitive variant (Lige pH-S)^26^ (Fig 4a, Extended Data Fig 3d-e). Lige pH-S demonstrated a similar degree of off-rate driven pH sensitivity at pH 6.0 as Oma pH-S while maintaining a higher binding affinity at neutral pH.

**Fig. 4.**
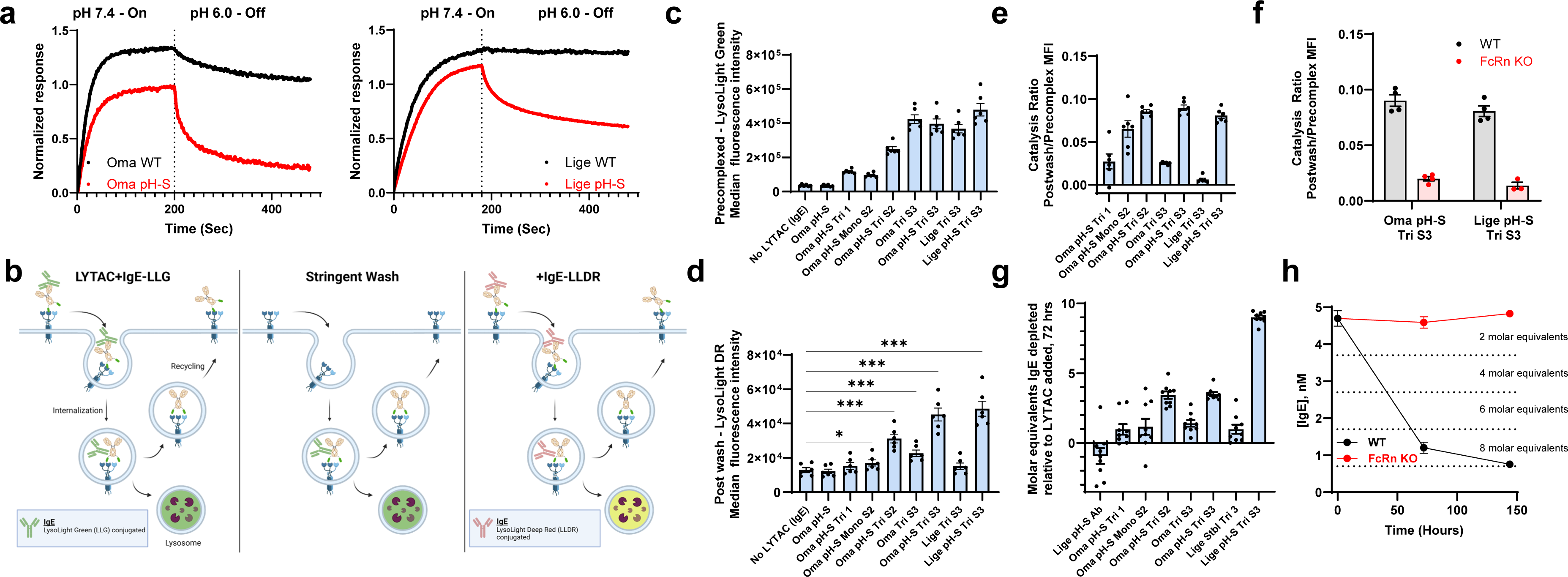
Design principles of catalytic LYTAC molecules. **(a)** BLI interferogram showing association of IgE to omalizumab (left) or ligelizumab (right) and associated pH-sensitive variants (pH-S, red) at pH 7.4, followed by disassociation at pH. 6.0. **(b)** Schematic describing catalytic uptake and degradation assay. LYTAC and LysoLight Green labelled IgE (IgE-LLG) were precomplexed before adding to HepG2 cells (first panel). After 2 hours of incubation, cells were stringently washed with low pH buffer on ice to arrest cycling and strip LYTAC off the cell surface (middle panel), followed by the addition of a second round of IgE conjugated to LysoLight Deep Red (IgE-LLDR). Using two separate fluorophore labelling schemes allows for differentiation between initial and catalytic degradation of target, before being read out by FACS. **(c-e)** FACS quantification of initial uptake and degradation (c), postwash catalytic degradation (d), and normalized catalytic degradation to initial LYTAC uptake/degradation (referred to as the catalytic ratio, e). **(f)** Catalytic ratio from FACS assay in HepG2 WT versus FcRn KO cells. **(g)** Direct measurement of catalytic depletion of IgE from media after 72 hour incubation with HepG2 cells. Molar equivalents of IgE depleted relative to LYTAC added calculated as [IgEinitial]-[IgEfinal]/[LYTACinitial]. **(h)** IgE depletion assay time course highlighting the Lige pH-S Tri S3 LYTAC in WT and FcRn KO HepG2 cells.

Introducing pH-sensitive target release into cycling LYTACs should, in principle, allow a single LYTAC molecule to internalize and deliver multiple target molecules for degradation, thereby realizing catalytic activity. Recycling antibodies similarly utilize pH-sensitive binding to a target protein to minimize target mediated antibody clearance, extend serum half-life and improve durability^27^. To determine whether the pH-sensitive LYTAC molecules are capable of catalytic target degradation, we developed two orthogonal in vitro assays. The first assay utilizes two different fluorescent cathepsin cleavable LysoLight probes (Green and Deep Red, LLG and LLDR, respectively) to evaluate multiple rounds of target internalization and degradation before and after washing out the LYTAC molecule (Fig 4b). A panel of LYTACs was evaluated, consisting of both original and pH-S variants of Oma and Lige, as well as a series of ASGPR ligands spanning different stabilities, avidities and affinities. In this assay, the pre-wash signal measures the uptake and degradation of pre-complexed LYTAC/IgE-LLG. We observed that the higher affinity ligands showed increased signal with no difference between pH-S vs. parental IgE binders (Fig 4c). The post-wash signal measures the ability of an internalized LYTAC:IgE-LLG complex to dissociate and recycle the LYTAC for additional round(s) of degradation as only recycled LYTAC will promote uptake/degradation of the differentially labelled IgE (IgE-LLDR). In contrast to the pre-wash signal, we observed that the pH-S IgE binders consistently outperformed non-pH sensitive binders (Fig 4d). A relative measure of catalytic activity can be estimated by the ratio of the pre/post wash signals. We observed the highest catalytic activity with LYTACs incorporating both stabilized ligands (Tri 1 vs. Tri S2, S3) and pH-S IgE binding (Oma vs. Oma pH-S) with no difference between the two IgE binders in this assay (Oma pH-S vs. Lige pH-S) (Fig 4e). Consistent with the previous cycling studies (Fig 3), catalytic activity was diminished in the absence of FcRn (Fig 4f).

To confirm that the stabilized, pH-sensitive LYTACs were capable of catalytic activity, an orthogonal in vitro depletion assay was developed to quantify the amount of IgE that can be degraded by each LYTAC molecule. HepG2 cells were incubated with 0.5 nM LYTAC and 10-fold molar excess IgE (5 nM) for 3 days. IgE concentrations at the initial timepoint and after 3 days were measured. Consistent with the FACS catalysis assay results, only LYTACs with stabilized ligands and pH-sensitive binding were capable of superstoichiometric (>2-fold) depletion of IgE from media (Fig 4g). The Lige pH-S LYTAC outperformed the Oma pH-S variant and approached the theoretical maximum depletion of 10 molar equivalents of IgE (9.0) for the assay (Fig 4g). The increased activity of Lige pH-S in this assay can likely be attributed to the higher affinity of Lige to IgE (Extended Data Fig 3d-e) and the low concentrations of IgE used in the depletion protocol^28^. As observed in the catalysis assay, FcRn was also required for depletion, with no quantifiable depletion of IgE in its absence after 6 days (Fig 4h). Taken together, using two independent assessments, these results demonstrate that ASGPR ligand stabilization, pH-sensitive target release and FcRn mediated cycling are the essential components conferring cycling LYTAC catalytic degradation of IgE.

### CataLYTACs Drive Deep and Durable Depletion of a Soluble Target Protein in vivo

To evaluate performance of cataLYTACs in a setting with constant production of a soluble target protein, we developed a xenograft mouse model utilizing hIgE-secreting U266B1 myeloma cells (U266). U266 cells were implanted in the flank of NOD-scid IL2Rgnull mice and allowed to expand over time, resulting in constantly increasing production and serum accumulation of hIgE over the course of the study (Fig 5a). Following a pre-dose determination of hIgE serum concentration, LYTACs were subcutaneously administered at 10 mg/kg and depth of hIgE depletion was quantified as a percentage of starting IgE concentration at 96h (Fig 5b). LYTACs incorporating pH-sensitive target binding and stabilized ASGPR binders outperformed counterparts lacking those modifications in the mouse constant production model (Fig 5b) and were well correlated with the in vitro FACS catalysis results (Fig 5c). We next sought to determine duration of activity provided by cataLYTACs in this model. Both **Lige pH-S** and **Oma pH-S Tri S3** LYTACs demonstrated deep and sustained depletion of hIgE with apparent pharmacodynamic activity lasting at least 2 weeks compared to a vehicle control (Fig 5d).

**Fig. 5.**
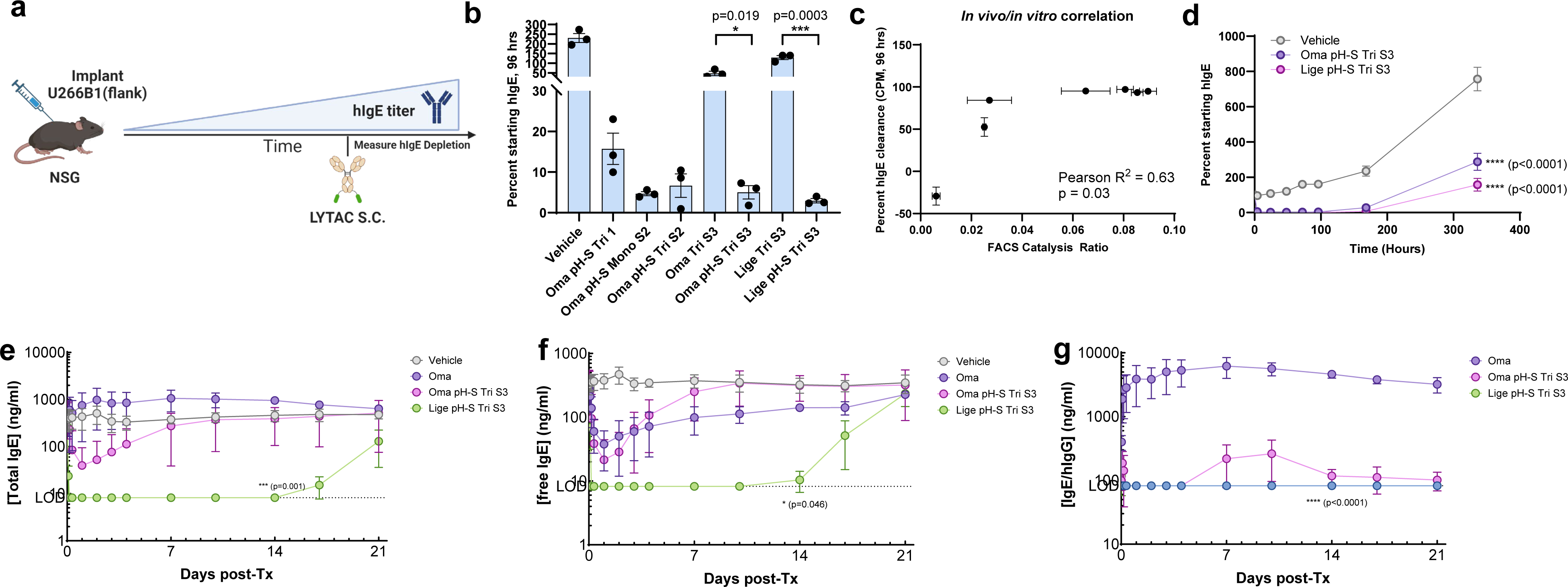
Catalytic LYTACs drive deep and durable *in vivo* depletion of soluble target protein in mouse and NHP. **(a)** Schematic of mouse model with constant production of hIgE via implantation of U266B1 myeloma cells. **(b)** Serum concentration of hIgEin mouse constant production model (“mouse CPM”) 96 hours after dosing of indicated test articles S.C. at 10mg/kg (n=3/group). Statistical analysis: Unpaired t test. **(c)** Correlation analysis of*in vitro* activity from Figure 4e and performance in mouse CPM depicted in Figure 5b. **(d)** Serum concentration of hIgE as percent of starting concentration over time in mouse CPM following S.C. administration of catalytic LYTACs at 10mg/kg (n=3/group).Data compiled from from two independent experiments. Statistical analysis: 2way ANOVA with Tukey’s multiple comparisons test, significance calculated compared to Vehicle group at 336hr timepoint. **(e)** Serum concentration of total IgE in cynomolgus macaques after single dose of indicated test article S.C. at 5 mg/kg (n=3/group). Statistical analysis: 2way ANOVA with Tukey’s multiple comparisons test, significance calculated compared to Vehicle group at 14d timepoint. **(f)** Serum concentrationof freeIgE in cynomolgus macaques after single dose of indicated test article S.C. at 5 mg/kg (n=3/group). **(g)** Serum concentrationof IgE/hIgG complexin cynomolgus macaques after single dose of indicated test article S.C. at 5 mg/kg (n=3/group). Statistical analysis for (f) and (g): 2way ANOVA with Tukey’s multiple comparisons test, significance calculated compared to Oma group at 14d timepoint.

The remarkable depth of depletion and duration of action by cataLYTACs in mice motivated us to explore whether these results translated to non-human primates (NHP)^29^. The **Lige pH-S Tri S3**, **Oma pH-S Tri S3** LYTACs are cross-reactive with cynomolgus macaque IgE and ASGPR, enabling a pharmacodynamics study that 1) does not require exogenous introduction of hIgE and inherently provides target resynthesis at a more physiological rate than the myeloma-based mouse model. Naïve cynomolgus macaques were pre-screened for IgE serum concentration and then subcutaneously dosed with 5mg/kg of either **Lige pH-S Tri S3**, **Oma pH-S Tri S3** or parental omalizumab. Both LYTACs drove an immediate reduction of total and free IgE with minimal formation of hIgG/IgE complexes, consistent with a degradation mechanism of action (Fig. 5e-g). **Lige pH-S Tri S3** dramatically outperformed the **Oma pH-S Tri S3** LYTAC in this model, providing more evidence that target binding is a critical driver of catalytic LYTAC activity. Omalizumab caused an increase in total IgE and hIgG/hIgE complexes, while reducing free IgE, as expected for a blocking antibody and consistent with clinical reports (Fig. 5g)^30^. **Lige pH-S Tri S3** drove a more rapid and deeper reduction of free IgE, a key measurement of efficacy, than Oma, dropping below the limit of detection within 4 hours of dosing (Fig. 5e-f)^31^. Importantly, this improved reduction in free IgE compared to Oma at an identical dose was durable, persisting for at least 2 weeks after administration (Fig 5f). This result not only demonstrates the potency and duration of action achieved by cataLYTACs, but also provides direct evidence that a catalytic LYTAC can outperform a clinically approved blocking antibody for neutralization of a soluble target protein.

## Discussion

Dose level and feasibility for occupancy-based antibody therapeutics are strongly dependent on several system specific parameters necessary to establish a therapeutic equilibrium state. Higher expression levels of target, faster target turnover, and stronger competing native interactions of a target all necessitate higher dose to achieve desired suppression of free target^32^. In addition, target and antibody pharmacokinetics can be interdependent, as in the case of target mediated drug disposition (TMDD) and antibody mediated target accumulation, requiring higher and/or more frequent dosing^33^. In contrast, event-driven degradation creates a non-equilibrium system that should relieve dependence on these parameters, particularly in the case where the modality can act in a catalytic fashion. Additionally, a degradation mechanism of action can be advantageous as it will impart maximal pharmacology associated with a pathogenic protein target which may not be achieved by blocking a single epitope with a standard monoclonal antibody.

These studies report the development of catalytically active ASGPR-directed cycling LYTACs that promote deep and sustained depletion of IgE in mouse and non-human primate model organisms. These cataLYTACs employ three key design features: 1) stabilized ASGPR ligands to mediate internalization, 2) pH-sensitive binding to facilitate target release in the endosome and FcRn-mediated retrieval of LYTACs from the degradation pathway. Our results demonstrate that all these design features are necessary for superstoichiometric depletion in vitro and optimal clearance and durability in vivo (Figs 4e-h, 5b). As demonstrated here for IgE in an NHP model (Fig 5e-g), the cataLYTAC provides substantially superior depth and duration of suppression of free target at equivalent dose to blocking antibody omalizumab. These data strongly support our hypothesis that an event-driven MOA can differentiate from occupancy-based MOAs. In addition, the engineered fate separation of the cataLYTAC (recycled) from IgE (degraded) eliminates antibody-mediated IgE accumulation observed with omalizumab that necessitates high dose for free IgE suppression.

The co-expression of ASGPR and FcRn in hepatocytes forms the biological basis for the activity observed with cataLYTACs and differentiates from what may be limiting for other approaches. For example, recycling/sweeping antibodies use FcRn and pH-sensitive binding to remove pathogenic proteins from circulation but do not exploit an efficient receptor-mediated endocytosis pathway to promote rapid and efficient internalization^27,34^. This approach, to our knowledge, has not been reported to result in deep and sustained depletion of an abundant (e.g. >10 ng/ml) constantly produced target from circulation, likely due to internalization limitations by receptor-independent pinocytosis or avidity-dependent internalization through immune complex cross-linking of Fc gamma receptors. ^35-37^. Recently, an FcRn-independent cycling approach utilizing transferrin receptor for membrane protein target degradation was reported^38^. While this receptor-mediated recycling approach is mechanistically simpler than the FcRn-dependent mechanism described here, the half-life of the therapeutic may be limited by the half-life of the receptor (<48 hrs) rather than exploiting the half-life extension imparted by FcRn recycling for antibodies and human serum albumin (∼3 weeks)^39,40^.

Finally, we anticipate cataLYTAC design features can be tuned to achieve maximum event-driven pharmacology, e.g.: 1) ASGPR ligand affinity at pH 7.4 and pH 6.0 for optimal receptor binding; 2) pH-sensitivity and dissociation rates for optimized endosomal target release; and 3) interaction with FcRn via addition of enhancing mutations for improved recycling. Since in vitro catalysis and in vivo clearance is well correlated, further optimization along these three axes is an experimentally viable path to produce cataLYTACs with favorable dosing regimens to treat a wide variety of diseases driven by soluble secreted proteins.

## Methods

### Chemical Synthesis

Structures and synthetic details are provided in Extended Data Fig 5 and the Supplementary Methods.

### Fluorescent Polarization Competitive Assay

ASGPR binding of GalNAc ligands was measured in black 384-well flat bottom plates (Greiner Bio, 07-000-890) using a fluorescence polarization competitive assay. A fluorescent probe consisting of a tri-GalNAc conjugated with Cy5 dye (**18**, Extended Data Fig 5) was custom synthesized. Ligands tested were resuspended in DMSO and 3-fold serial dilutions were made at 100x final concentrations. Binding reactions were conducted in 50 µl final volume in 20 mM HEPES (pH 7.5), 50 mM NaCl, 5 mM CaCl_2,_ 0.015% Triton X-100, and 1% DMSO with 100 nM recombinant ASGPR (Acro Biosystems, GS1-H82Q3) and 1 nM Cy5 probe. Fluorescence polarization was measured using λ_ex_ = 620 nm, λ_em_ = 688 nm on an Envision plate reader (Perkin Elmer) after 2 hr incubation time at 25 °C. Dose responses were conducted in triplicate and normalized to the response with DMSO (100%) and 1 µM reference compound **6** (0%) on each plate. IC_50_ values were determined by fitting to 4-parameter non-linear curves in GraphPad Prism.

### Antibody Expression and Purification

All anti-IgE antibodies were cloned into a pTT5 vector (ThermoFisher Scientific) with a hIgK signal peptide and human IgG1 Fc backbone containing a LALA mutation for effector-attenuation, a YTE mutation for enhanced FcRn binding, and a L443C mutation for site-specific conjugation. The antibodies were transiently expressed in HEK293 according to the manufacturer’s protocol (ThermoFisher Scientific). Antibodies were purified on FLPC using a two-step purification process. First, antibodies were purified from cell culture medium using Protein A affinity chromatography according to the manufacture’s protocol (MabSelect SuRe; Cytiva, 17543801). Antibodies were then polished with an additional purification step to remove remaining contaminates and high molecular weight species by cation exchange chromatography (HiTrap SP HP; Cytiva, 17115201). Antibody purity was measured with HPLC size exclusion chromatography (Sepax ZENIX-C SEC-300) with 2X Phosphate buffered saline as the mobile phase. The identity of the antibodies was confirmed by mass spectral (MS) characterization using Xevo-TOF-MS.

### Antibody-GalNAc Conjugation

Following two-step purification, antibodies were buffer exchanged into HBSE, pH 7.2 buffer (25 mM HEPES, 150 mM NaCl, 1 mM EDTA) using Zeba Desalting Columns (ThermoFisher Scientific, A57764) and diluted to 5-10 mg/mL. To remove cysteine capping on the L443C site, 20 molar equivalents of tris(2-carboxylethyl) phosphine (TCEP; ThermoFisher Scientific, 77720) was added to the antibodies and the mixture was incubated at 25°C for 2 hours. TCEP was removed from the reaction mixture using Zeba Desalting Columns equilibrated with fresh HBSE, pH 7.2 buffer. Removal of cysteine capping the L443C site was confirmed with Xevo-TOF-MS. An 80 mM stock solution of L-dehydroascorbic acid (DHAA; Sigma Aldrich, 261556) was prepared fresh in HBSE, pH 7.2 buffer. To re-oxidize the antibodies, 40 molar equivalents of DHAA was added to the antibodies and incubated for 2 hours at 25°C. Re-oxidation was confirmed by SDS-PAGE. Next, 5 molar equivalents of maleimide-GalNAc were added to the de-capped antibodies and reacted at 37°C for 2 hours. LYTAC conjugates tested in mice with constant production of hIgE or cynomolgus macaques underwent an additional step to stabilize the linkage by thiosuccinimide hydrolysis. For these conjugates, the pH of the reaction mixture was adjusted to 8-8.3 using 1M Borate, pH 9 and incubated overnight at 37°C. Unreacted GalNAc and high molecular weight species were removed by FPLC using size exclusion chromatography (Superdex 200 Increase 10/300 GL; Cytiva, 28990944). The concentration of the LYTAC conjugates was measured using a Nanodrop. Conjugate purity was measured with HPLC size exclusion chromatography and Xevo-TOF-MS was used to estimate the linker/ligand:antibody ratio (Extended Data Fig 4).

### Cell line generation and maintenance

HepG2 cells were purchased from ATCC. ASGPR1/2 and FcRn knockout cell lines were generated by CRISPR-Cas9 genome editing. Guide RNAs were designed for ASGR1 (GGACGGGACGGACTACGAGA) and ASGR2 (TGATTGCCTGGACCTCCGTC) using the ALT-R design tool (Integrated DNA Technologies, IDT). CRISPR/Cas9 ribonuceleoproteins (RNPs) were assembled and electroporated into HepG2 cells using a Lonza Amaxa 4D nucleoporation system according to manufacturer’s instructions. Single cells were expanded by limited dilution plating and ASGPR1/2 double KO clones were identified by surface staining using ASGPR1 and ASGPR2 antibodies. FcRn (FGCRT) KO cells were generated using the Knockout Express platform (Synthego). Single cells were expanded and clones KO clones were confirmed by sequencing. All cell lines were maintained in Eagle’s minimal essential medium (EMEM) supplemented with 10% fetal bovine serum and 1x penicillin/streptomycin and incubated at 37°C with 5% CO_2_.

### LysoLight Deep Red IgE Cleavage Assay

Human IgE (Enzo BPD-DIA-HE1) was conjugated to the LysoLight Deep Red (LLDR) probe using standard protocol recommended by manufacturer (ThermoFisher Scientific, L36004), with two dye removal steps included to ensure sufficient removal of free dye, and stored at 4°C.

For cell-based assays, wild type or ASGPR1/2 KO HepG2 cells were plated at a density of 25k cells/per well in a 96-well flat bottom plate 48 hours before experiment (counted with ViCell XR Cell Viability Analyzer). Cells were ∼70% confluent on the day of experiment. The day of the experiment, 600 nM of LYTAC was precomplexed with equimolar concentration of LLDR-IgE in cell culture media for 30 minutes, before being serially diluted (Log3 dilutions). During this period cells were treated with Fc receptor blocking solution (Biolegend Human TruStain FcX, 1:100 dilution in fresh media). At the end of the precomplexing/dilution steps, the precomplexed LYTAC:LLDR-IgE dilutions were added to cells at equivalent volumes as the existing media with Fc receptor blocking solution, for a final concentration range of 1.2-300 nM. Two separate controls with LLDR-IgE without LYTAC and with cathepsin inhibitor (Aloxistatin/E64d, Selleck Chemical S7393) added with the Fc receptor block (final concentration of 5 μM during assay) were included to account for high background with the LLDR-IgE probe and to confirm that fluorescence signal was due to cleavage by cathepsins in the lysosome. After incubating for two hours in cell incubator, cells were washed with DPBS and disassociated with 0.25% trypsin/EDTA disassociation solution for 10 minutes before transferring to conical bottom 96-well plates. Cells were centrifuged, washed with FACS buffer (with BSA, Rockland Immunochemicals), centrifuged again and resuspended in FACS buffer for analysis on Agilent NovoCyte Advanteon. For one replicate, an additional step between washes was included where cells were incubated with Zombie Aqua^TM^ dye (Biolegend, 1:500) in DPBS on ice for 20 minutes before being centrifuged and resuspended for flow cytometry analysis; using a live/dead dye confirmed that gating on size alone was sufficient to include the live cell population for HepG2 cells. For each well, 3,000 live cells were collected and analyzed.

### GalNAc catabolism assay

A mixture of LYTAC molecules (25 nM final concentration), 1x catabolic buffer (from 10x catabolic buffer, XenoTech, LLC), and rat liver tritosomes (diluted 1:20, from BioIVT) was made on ice and volumed up to 50 μl with cold MilliQ H_2_O. Half the reaction was flash frozen and stored at -80°C for analysis of the initial timepoint, while the other half was incubated at 37 degrees for 3 hours before flash freezing and storing at -80°C. ELISAs were then performed to quantify IgG and GalNAc in the initial and three-hour timepoint samples. 96-well immuno plates were coated with 1 μg/ml of α-HuIgG (Invitrogen A18827, IgG ELISA) or 2 μg/ml α-HuIgG (SouthernBioTech 2049-01, GalNAc ELISA) in PBS at 4°C overnight. Plates were washed three times with DPBS with 0.05% Tween-20 and blocked with PBST with 1% w/v BSA for one hour. Samples were then thawed on ice, diluted in PBST with 1% w/v BSA and 0.05% Tween-20 (IgG dilution buffer) or TBST with 1% w/v BSA and 5 mM Ca^2+^ (GalNAc dilution buffer). For the IgG ELISA, unconjugated Oma was used as a standard for all LYTACs; for the GalNAc ELISA, separate standard curves were generated for each unique GalNAc ligand to account for differences in GalNAc affinity to the ASGPR detection reagent. Samples and standards were incubated on the blocked plates for one hour before washing three times with wash buffer (PBST for IgG ELISA, TBST with 5 mM Ca^2+^ for GalNAc ELISA), followed by addition of detection reagent (α-human IgG-HRP, Sigma SAB3701283 for IgG ELISA, and 0.5 μg/ml biotinylated ASGPR1 from Acro Biosystems for GalNAc ELISA) for one hour. For the GalNAc ELISA plates, a secondary detection reagent was added for one hour (Streptavidin HRP, Pierce 21130, 1:3000 in GalNAc dilution buffer). Detection of HRP reagents for both ELISAs was done using 3,3’,5,5’-Tetramethylbenzidine (Sigma-Aldrich T0440) followed by quenching with 0.18 M sulfuric acid. Plates were read on a Perkin Elmer EnVision 2105 plate reader.

### LYTAC cycling assays

Human IgE (Enzo BPD-DIA-HE1) and LYTACs were conjugated to Alexa Fluor 647 and Alexa Fluor 488, respectively, according to manufacturer’s instructions (Invitrogen Protein Labeling Kits, A10235 and A20173), using half the amount of recommended dye for conjugation.

For LYTAC cycling assays described in Fig. 3b and S2a-e, HepG2 cells were plated at a density of 300,000 cells per well in 12 well plates 48 hours before the experiment. On the day of the experiment, 100 nM of Alexa Fluor 488 labeled LYTAC molecules in cell culture media was added to cells for one hour in incubator. Afterwards, cells were placed on ice and washed three times with ice cold buffers, once with low pH/calcium wash buffer (20 mM sodium acetate, 150 mM NaCl, pH 5.2), once with EBSS adjusted to pH 2.5, and once with DPBS (30-45 seconds for each step). After wash steps, 200 μl of cold 0.25% trypsin/EDTA was added to each well and cell disassociation was allowed to proceed for 10 minutes on ice before adding 800 μl of prewarmed cell culture media with 1 μg/ml Alexa Fluor 647 labelled α-HuIgG (Invitrogen A48279). For experiments with GalNAc competitor during cycling steps, 1 μM of Tri S3 analogue used in fluorescence polarization assays was also added. Cells were removed from plate and put in Eppendorf tubes and placed in cell incubator. Every 15-30 minutes for 90 minutes, 150 μl cell aliquots were removed from tubes and placed in fresh Eppendorf tubes on ice. Cells were transferred to cold conical bottom 96-well plate and washed and analyzed by flow cytometry as described above (LysoLight Deep Red IgE Cleavage Assay).

For cycling assays described in Fig. 3c-d and S2f-h, HepG2 cells were plated at a density of 25,000 cells per well in 96-well plates 48 hours before the experiment. The day of the experiment, serial dilutions were made with the Alexa Fluor 488-labeled Oma Tri S2 and efgartigimod, such that the LYTAC concentration was constant at 50 nM but the efgartigimod concentration was serially diluted 1:2 for a concentration series ranging from 3-1,000 nM, with a final LYTAC only stock included as a control. The dilution series was incubated with cells for 90 minutes to internalize in a cell incubator. Afterwards, cells were placed on ice and washed with cold low pH buffers and DPBS as described in the previous section, before fresh media containing 100 nM Alexa Fluor 647 conjugated IgE and identical efgartigimod dilution series was added to cells. Cells were incubated in incubator for four hours before washing, detaching, and preparing for flow cytometry analysis as described in previous sections.

For imaging-based cycling studies, HepG2 cells were plated at a density of 20,000 cells per well in Ibidi 18-well chambered slides, pretreated with poly-L-ornithine. 48 hours later, fresh cell culture media with 100 nM Alexa Fluor 488 labeled-Oma Tri S2 was added. Two hours later, cells were washed twice with cold DPBS, followed by addition of fresh cell culture media with 500 nM Alexa Fluor 647-labeled IgE. After four hours, cells were washed with PBS, fixed with 4% formaldehyde Fixative Solution in PBS (Invitrogen FB002), washed with PBS, and permeabilized with 0.1% Triton X100. Cells were washed with PBS before blocking for one hour with PBS with 5% goat serum. Afterwards, primary antibody (α-LAMP1, Cell Signaling 9091, diluted 1:400) in PBS with goat serum was added and let incubate at 4°C overnight. The following day, primary antibody was removed, washed 3 times with PBS, and secondary antibody (α-Rabbit IgG conjugated to Alexa Fluor 555, Cell Signaling 4413, diluted 1:1,000) in PBS with 5% donkey serum and 0.05% Tween 20 was added at room temperature for three hours. Cells were then washed 3 times with PBS with Hoescht 33342 (1:10,000) before adding two drops of Prolong Glass^TM^ antifade reagent to each well. Slides were stored at 4°C before being imaged on a Leica Stellaris 5 confocal microscope. Two to three separate frames were imaged per condition, and Leica LAS X colocalization software was used to quantify colocalization.

### LysoLight Deep Red LYTAC lysosomal cleavage assay

Oma Tri S2 was conjugated to LysoLight Deep Red (LLDR) as previously described. HepG2 cells were plated at a density of 25,000 cells per well 48 hours before the experiment. Day of the experiment, stocks of 25 nM of Oma Tri S2-LLDR were prepared in cell culture media, with or without 1 μM efgartigimod. Log2 serial dilutions were made to vary efgartigimod concentration but keep LYTAC-LLDR concentration consistent, as described for cycling assays. Dilution series was added to cells, let incubate in incubator for four hours, and then washed and prepared for flow cytometry analysis as described in previous sections.

### Generation of pH-sensitive IgE binders

Initial screening for pH-sensitive Omalizumab variants was performed using BLI on a Gator Prime instrument. Anti-human IgG Fc Gen II probes (Gator 160024) were equilibrated and baselined in BLI buffer (20 mM citric acid/phosphate, 150 mM NaCl, 0.05% Tween-20, pH 7.4 or 6.0) for 120 seconds before loading the probes with 1 μg/ml Omalizumab single His variants for 240 seconds. Following another 120 second baseline step, probes were immersed in varying concentrations of human IgE solutions (BioRad HCA171, concentrations from 0.3-20 nM with Log2 dilutions) for 200 seconds (association) before being moved to buffer only solution for 300 seconds (disassociation). Data was globally fit using Gator analysis software. For experiments to characterize pH-dependent disassociation in the pH-S variants of Oma and Lige, experiments were performed as described at a single IgE concentration (20 nM for Omalizumab variants and 10 nM for Ligelizumab variants), but loading of antibodies and association to IgE were performed in pH 7.4 buffer, and disassociation step was performed at pH 6.0.

IgE binding kinetics for combination histidine mutants of Omalizumab and Ligelizumab were measured at pH 7.4 and pH 6.0 using surface plasmon resonance (SPR). SPR was performed on Biacore T200 instrument. A monoclonal anti-human IgG antibody (Thermo Fisher Scientific, A55735) was conjugated to both reference and active flow cells of Series S CM5 chips using amine coupling. Recombinant human IgE (BioRad HCA171G) binding was performed in 20mM citric acid phosphate at either pH 7.4 or 6.0, 150mM NaCl, 0.01% Tween-20. Omalizumab or Ligelizumab variants were immobilized on the anti-IgG chip, paired with reference flow cells where no IgG was immobilized. IgE was flowed over at 7 increasing concentrations in a multi-cycle kinetic assay, with 120 second association and 600s final dissociation. All flow cells of the chip were regenerated between variants with 10mM pH 2.0 glycine buffer for subsequent immobilization and binding cycles. Binding data was fit to either bivalent analyte or one-to-one binding models with Biacore T200 Evaluation Software 3.2.1 as indicated.

### FACS two-color catalysis assay

HepG2 cells were plated at a density of 25k cells/per well in a 96-well flat bottom plate 48 hours before the experiment. Two wells per condition were plated for replicates. The day of experiment, the IgE conjugated to LysoLight Green (ThermoFisher Scientific, L36006) was precomplexed with the LYTAC molecule at equal stoichiometries (100 nM) in cell culture media for 20 minutes; media was then removed from cells and replaced with fresh media with precomplexed LYTAC/IgE. Cells were incubated with LYTAC:IgE in cell incubator for 2 hours. Afterwards, cells were placed on ice and washed three times with ice cold buffers, once with low pH/calcium wash buffer (20 mM sodium acetate, 150 mM NaCl, pH 5.2), once with EBSS adjusted to pH 2.5, and once with DPBS. For each washing step, buffer remained on cells for 30-45 seconds. Once washing was complete, fresh media with 50 nM IgE conjugated to LysoLight Deep Red was added. Cells were incubated in cell incubator for 4 hours. Media was then removed from cells, and cells were treated for flow cytometry analysis as described above (LysoLight Deep Red IgE Cleavage Assay).

### In vitro IgE Depletion Assay

HepG2 cells were plated at a density of 75,000 cells per well in the inner 24 wells of a 48 well plate. All outer wells of the plates were filled with 1 ml of DPBS to minimize evaporation over the course of the experiment. 24 hours later, media was replaced with 100 μl of cell culture media with 5 nM human IgE (Bio-Rad, HCA171) and 0.5 nM LYTAC. A portion of the LYTAC:IgE mix was saved and frozen at -20°C to accurately quantify IgE starting concentrations before depletion. Cells were incubated with LYTAC for 72 hours before plates were spun down to pellet any cell debris and media was collected and frozen at -20°C. For experiments with FcRn KO cells, media aliquots were collected at 72 and 144 hours.

IgE concentrations in starting and cell treated samples were analyzed using Meso Scale Discovery R-PLEX Human IgE Assay, as per manufacturer’s instructions.

### Mouse Models with Human IgE

All animal studies were conducted in an AAALAC accredited facility under IACUC-approved protocol.

*Pharmacodynamics:* Female C57BL/6J mice (The Jackson Laboratory; strain #000664) were dosed I.V. with 1mg/kg of human IgE (Bio-Rad, HCA171), followed by 5mg/kg test article I.V. the next day. Serum samples were collected prior to administration of test articles and at 1 hour, 4 hours, and 24 hours after dosing. N=3 mice per experimental group. Serum hIgE concentrations were measured by Meso Scale Discovery R-PLEX Human IgE Assay.

*Delayed-challenge pharmacodynamics:* Female C57BL/6J micewere dosed I.V. with 5mg/kg test articles, followed by 1 mg/kg of human IgE (Bio-Rad, HCA171) I.V. the next day. Serum samples were collected at 5 minutes, 1 hour, 4 hours, and 24 hours after administration of hIgE. N=3 mice per experimental group. Serum hIgE concentrations were measured by Meso Scale Discovery R-PLEX Human IgE Assay.

*Constant production:* 5 x 10^6^ U266B1 myeloma cells (TIB-196, ATCC) were implanted subcutaneously into the flank of female NOD-scid IL2Rgamma-null mice (The Jackson Laboratory strain #005557). Tumor growth and hIgE titers (via Meso Scale Discovery R-PLEX Human IgE Assay) were monitored to ensure successful myeloma implantation. Prior to administration of test articles, mice were divided into experimental cohorts with normalized levels of hIgE. LYTACs and controls were then dosed subcutaneously into the scruff of the neck at 10mg/kg. N=3 mice per experimental group. Blood was collected at indicated timepoints, with serum subsequently tested for depletion of hIgE using Meso Scale Discovery R-PLEX Human IgE Assay.

### Non-human Primate (NHP) Pharmacodynamics Study

The NHP study was conducted by Wuxi AppTec in accordance with the WuXi IACUC standard animal procedures along with the IACUC guidelines that are in compliance with the Animal Welfare Act, and the Guide for the Care and Use of Laboratory Animals. Cynomolgus macaques were pre-screened for serum IgE concentration using 2 weeks prior to study initiation. Each test article was administered subcutaneously at 5mg/kg followed by post-dose blood collection at 30min, 1hr, 2hr, 4hr, 8hr, 1d, 2d, 3d, 4d, 7d, 10d, 14d, 17d, and 21d. The resulting serum fractions were then subjected to ELISA analysis for total IgE, free IgE and IgE/hIgG complex as described below.

### NHP Serum Analysis

#### MSD Detection Method for Free IgE in NHP serum

MultiArray HB 96-well plates (Meso Scale Discovery, L15XB-6) coated for 1 hour at RT with 1 µg/ml mouse anti-human IgE (Miltenyi, 130-039-127) were blocked with PBS-T 1% BSA for 1 hour at RT. Standard curve was generated using cyno IgE protein (Acro, IGE-C52H3) diluted in PBS-T 1% BSA serial diluted into the same buffer including equivalent IgE-free cyno serum. Serum samples were diluted 1:20 in PBS-T 1% BSA in duplicate and pipetted into the wells along with the standard and incubated for 1 hour at RT. 5 µl biotinylated FcER1 (Acro, FCA-H82E3) was added into the wells for 30 minutes at RT. After washing in PBS-T, 0.5 µg/ml sulfo-tag streptavidin (Meso Scale Discovery, R32AD-1) was added into the wells for 1 hour at RT. Plates were then read using 150 µl read buffer B (Meso Scale Discovery, R60AM-4) and electrochemiluminescence was measured on the MSD reader (Meso Scale Discovery, Meso Sector S 600MM). The concentration of free IgE in each sample was determined based on the standard curve on each plate. All incubation steps were done while shaking.

#### MSD Detection Method for Total IgE in NHP serum

MultiArray HB 96-well plates (Meso Scale Discovery, L15XB-6) coated for 1 hour at RT with 0.5 µg/ml mouse anti-human IgE (Miltenyi, 130-039-127) were blocked with PBS-T 1% BSA for 1 hour at RT. Standard curve was generated using cyno IgE protein (Acro, IGE-C52H3) diluted in PBS-T 1% BSA serial diluted into the same buffer including equivalent IgE-free cyno serum. Serum samples were diluted 1:20 in PBS-T 1% BSA in duplicate and pipetted into the wells along with the standard and incubated for 1 hour at RT. After washing in PBS-T, 1 µg/ml biotinylated anti-human IgE (BioRad, STAR147B) was added into the wells for 1 hour at RT. Plates were washed again and 0.5 µg/ml sulfo-tag streptavidin (Meso Scale Discovery, R32AD-1) was added into the wells for 1 hour at RT. Plates were then read using 150 µl read buffer B (Meso Scale Discovery, R60AM-4) and electrochemiluminescence was measured on the MSD reader (Meso Scale Discovery, Meso Sector S 600MM). The concentration of total IgE in each sample was determined based on the standard curve on each plate. All incubation steps were done while shaking.

#### MSD Detection Method for human IgG in NHP serum

Streptavidin-coated 96-well plates (Meso Scale Discovery, L15SA-1) coated for 1 hour at RT with 1 µg/ml biotinylated goat anti-human IgG (Southern Biotech, 2049-08) were blocked with PBS-T 1% BSA for 1 hour at RT. Serum samples diluted in PBS-T 1% BSA were serial diluted into the same buffer and pipetted into the wells and incubated for 1 hour at RT. After washing in PBS-T, 2 µg/ml sulfo-tag anti-human IgG (Meso Scale Discovery, D21APR-6) was added into the wells for 1 hour at RT. Plates were then read using 150 µl read buffer B (Meso Scale Discovery, R60AM-4) and electrochemiluminescence was measured on the MSD reader (Meso Scale Discovery, Meso Sector S 600MM). The concentration of human IgG in each sample was determined based on the standard curve on each plate. All incubation steps were done while shaking.

#### MSD Detection Method for IgE-IgG Complex (bound IgE) in NHP serum

MultiArray HB 96-well plates (Meso Scale Discovery, L15XB-6) coated for 1 hour at RT with 0.5 µg/ml mouse anti-human IgE (Miltenyi, 130-093-127) were blocked with PBS-T 1% BSA for 1 hour at RT. Standard curve was generated using molar equivalence of the test article IgG source and cynomolgus IgE protein (Acro Biosystems, IGE-C52H3) diluted in PBS-T 1% BSA serial diluted into the same buffer including equivalent IgE-free cyno serum. Serum samples were diluted 1:20 in PBS-T 1% BSA and pipetted into the wells along with the standard and incubated for 1 hour at RT. After washing in PBS-T, 1 µg/ml biotinylated goat anti-human IgG (Southern Biotech, 2049-08) was added into the wells for 1 hour at RT. Plates were washed again, and 0.5 µg/ml sulfo-tag streptavidin (Meso Scale Discovery, R32AD-1) was added into the wells for 1 hour at RT. Plates were then read using 150 µl read buffer B (Meso Scale Discovery, R60AM-4) and electrochemiluminescence was measured on the MSD reader (Meso Scale Discovery, Meso Sector S 600MM). The concentration of IgE-IgG complex in each sample was determined based on the standard curve on each plate. All incubation steps were done while shaking.

## Data Availability

All data is presented in the paper, Supplementary Information or is available upon reasonable request.

## Acknowledgements

We thank Carolyn Bertozzi, Steven Banik, Monther Abu-Ramaileh, Alanna Schepartz and Randy Schekman for helpful discussion of experiments related to this manuscript. We thank Sarah Totten and Gabriela De Los Santos for high-resolution mass spectrometry confirmation of the chemicals described herein. We thank Richard Laura, Carrie Boerem and Carly Cheung for their early-stage contributions to this work. Schematics were generated using BioRender software. All work was funded by Lycia Therapeutics Inc.

## Contributions

C.K., M.J.S. and N.A.L. designed the research. C.K. and M.J.S. contributed equally to this work and are listed alphabetically. C.K. and M.J.S. designed and executed the in vitro cycling, catalysis and mechanism of action studies. N.L. designed and supervised the in vivo studies. R.M.L. designed the conjugation strategy, generated the LYTACs and performed quality control analysis. T.C. designed, supervised and executed the chemical syntheses. C.L.C. and R.K. developed the assays and performed the in vivo bioanalytical work. I.E.J. designed and executed the studies identifying the pH-sensitive omalizumab variants. S.G. designed and performed in vitro ASGPR binding assays. K.T.A. designed and supervised cloning/expression of the omalizumab and ligelizumab variants described. K.H.D. and D.L. purified and performed quality control analysis of omalizumab and ligelizumab variants. C.K.F. and T.T. performed the in vivo mouse studies. D.J.H. and S.R. performed the chemical syntheses. R.Y. designed cycling LYTACs and performed cloning/expression of omalizumab and ligelizumab. R.W.H. and C.L. developed the lysolight probes, provided advanced access and gave input on application to in vitro studies. D.A.L. designed cycling LYTAC framework, supervised LYTAC and other reagent generation. S.M.M. supervised NHP and other in vivo studies. J.S.I. designed and supervised in vivo bioanalytical studies, in vitro binding studies and pH-sensitive binder screening. J.G.L. and E.D.T. designed stabilized ASGPR ligands and supervised syntheses. S.T.S. supervised research related to ligand stabilization and selection of IgE as the model system to demonstrate cataLYTAC activity. S.T.S. also edited and revised the manuscript. C.K., N.A.L. and M.J.S. wrote the manuscript with input from all authors.

---

**Extended Data Fig 1.**
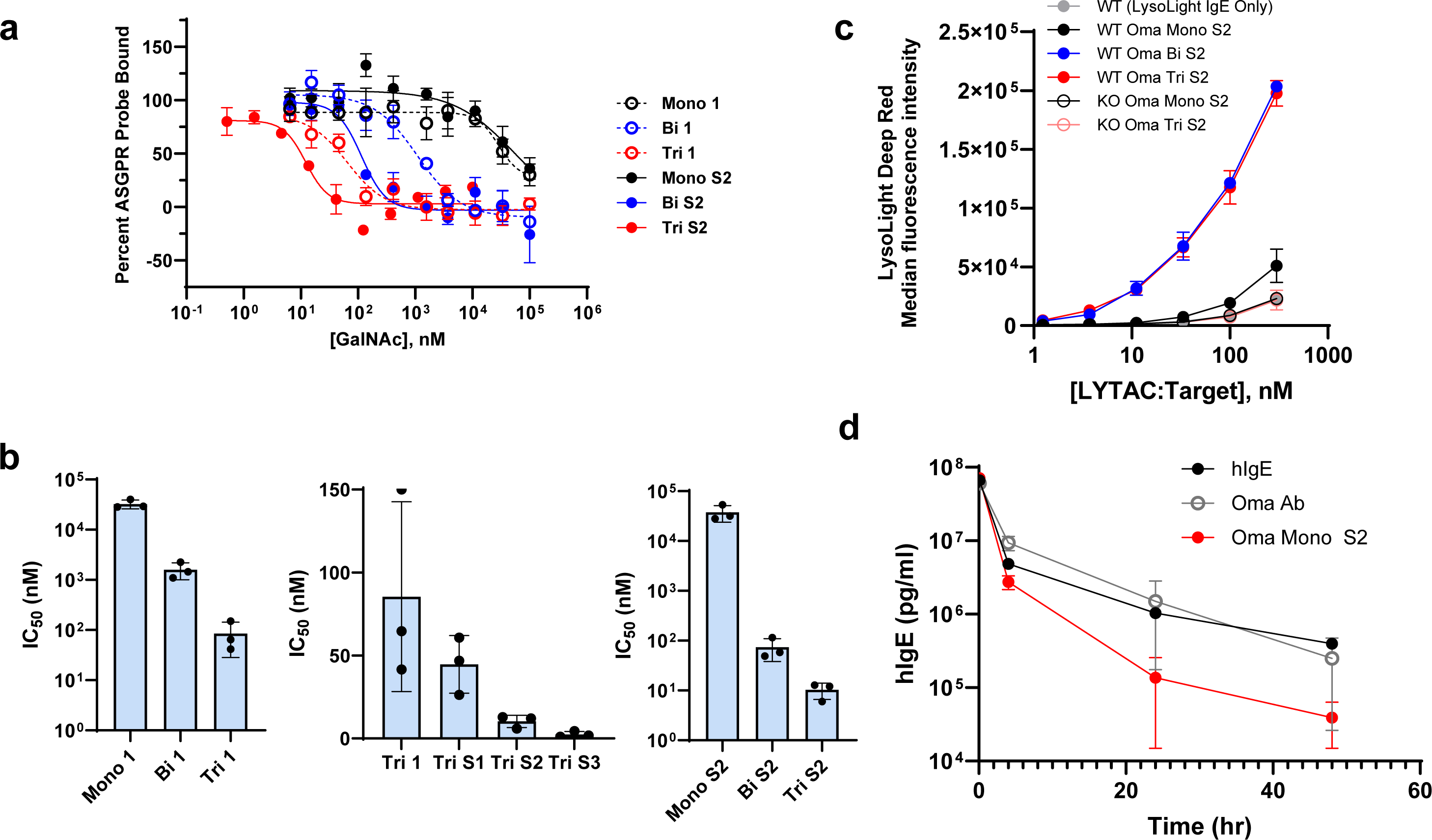
*In vitro* and*in vivo* characterization of engineered GalNAc ligands. **(a-b)** Fluorescence polarization competition assay of different valency GalNAc and stabilized GalNAc analog molecules with a fluorescently labelled TriGalNAc probe to recombinant ASGPR1.**(a)** Binding curves from GalNAc and GalNAc analog (S2) valency series. Data is representative from one biological replicate. **(b)** Summary of triplicate IC50values (from three biological replicates) of GalNAc valency series (left), trivalent GalNAc analogs (middle), and S2 GalNAc analog valency series (right).**(c)** Comparison of degradation efficiency of stabilized valency series LYTACs measured by *in vitro* cleavage of LysoLight Deep Red labelled-IgE by HepG2 WT or ASGPR1/2 KO cells. **(d)** In vivo delayed dosing experiment (analogous to Fig. 2e); 5.0 mg/kg of Oma/LYTAC was dosed intravenously 7 days prior to an intravenous 5.0 mg/kg dose of hIgE. Depletion of hIgEwas measured for 48 hours after hIgEdosing.

**Extended Data Fig 2.**
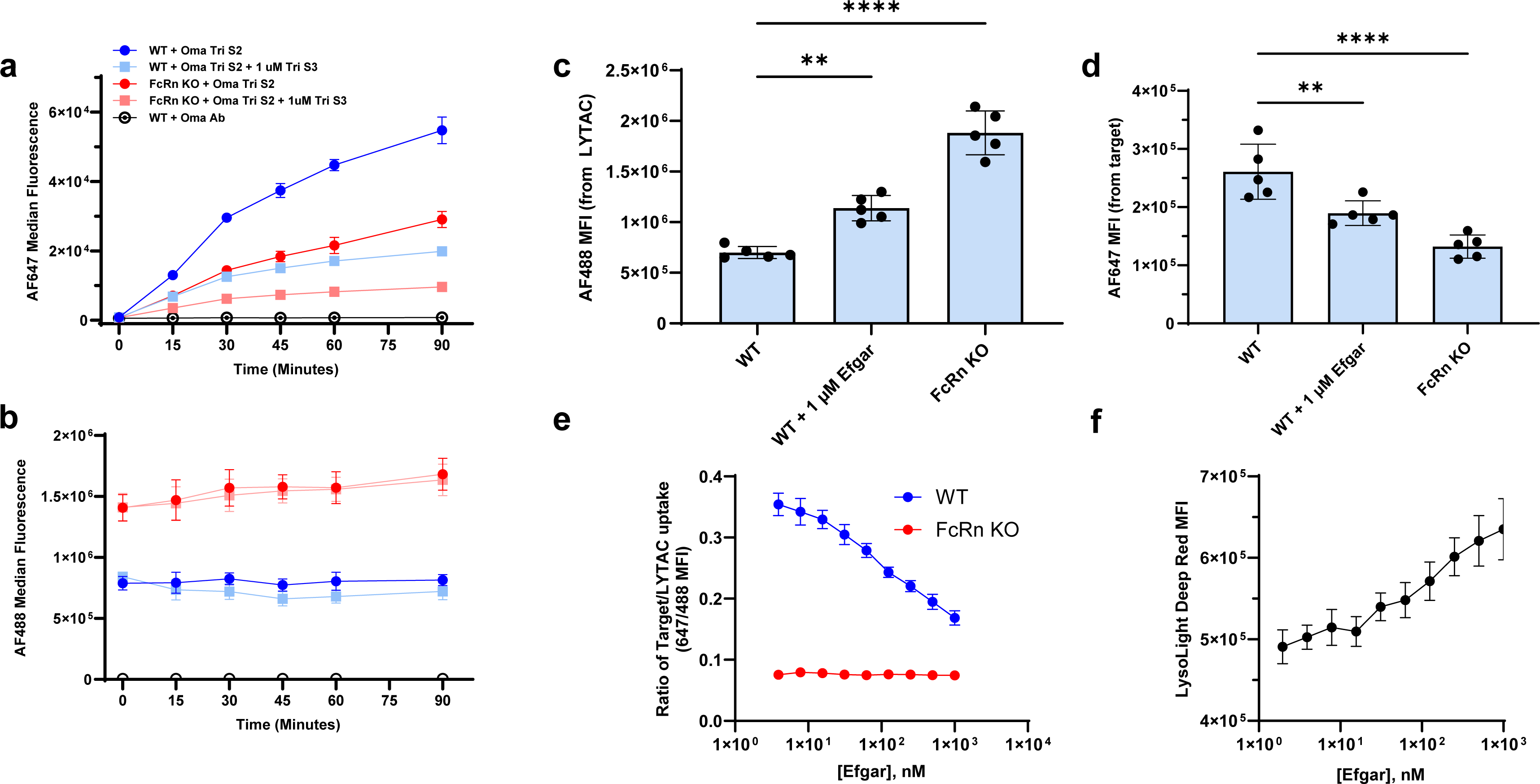
Elucidating the recycling pathways of cycling LYTACs. **(a)** Recycling signal (not normalized to LYTAC uptake) from Alexa Fluor 647 conjugated α-human IgG, corresponding to Fig. 3b. **(b)** LYTAC uptake in recycling assay measured by Alexa Fluor 488 signal from fluorescently conjugated LYTACs. **(c)** Oma Tri S2 (conjugated to Alexa Fluor 488) uptake in HepG2 WT cells compared to efgartigimod treated cells and FcRn KO (expanded from Fig. 3c). **(d)** Alexa Fluor 647 labelled IgE uptake in HepG2 WT cells compared to efgartigimod treated cells and FcRn KO (expanded from Fig. 3d). **(e)** Normalized ratio of IgE uptake to LYTAC uptake from Fig. 3c-d; FcRn KO cells are insensitive to treatment with efgartigimod **(f)** LYTAC catabolism measured by fluorescence of LysoLight Deep Red labelled Oma Tri S2 in the presence of FcRn inhibitor efgartigimod after coincubation for 4 hours.

**Extended Data Fig 3.**
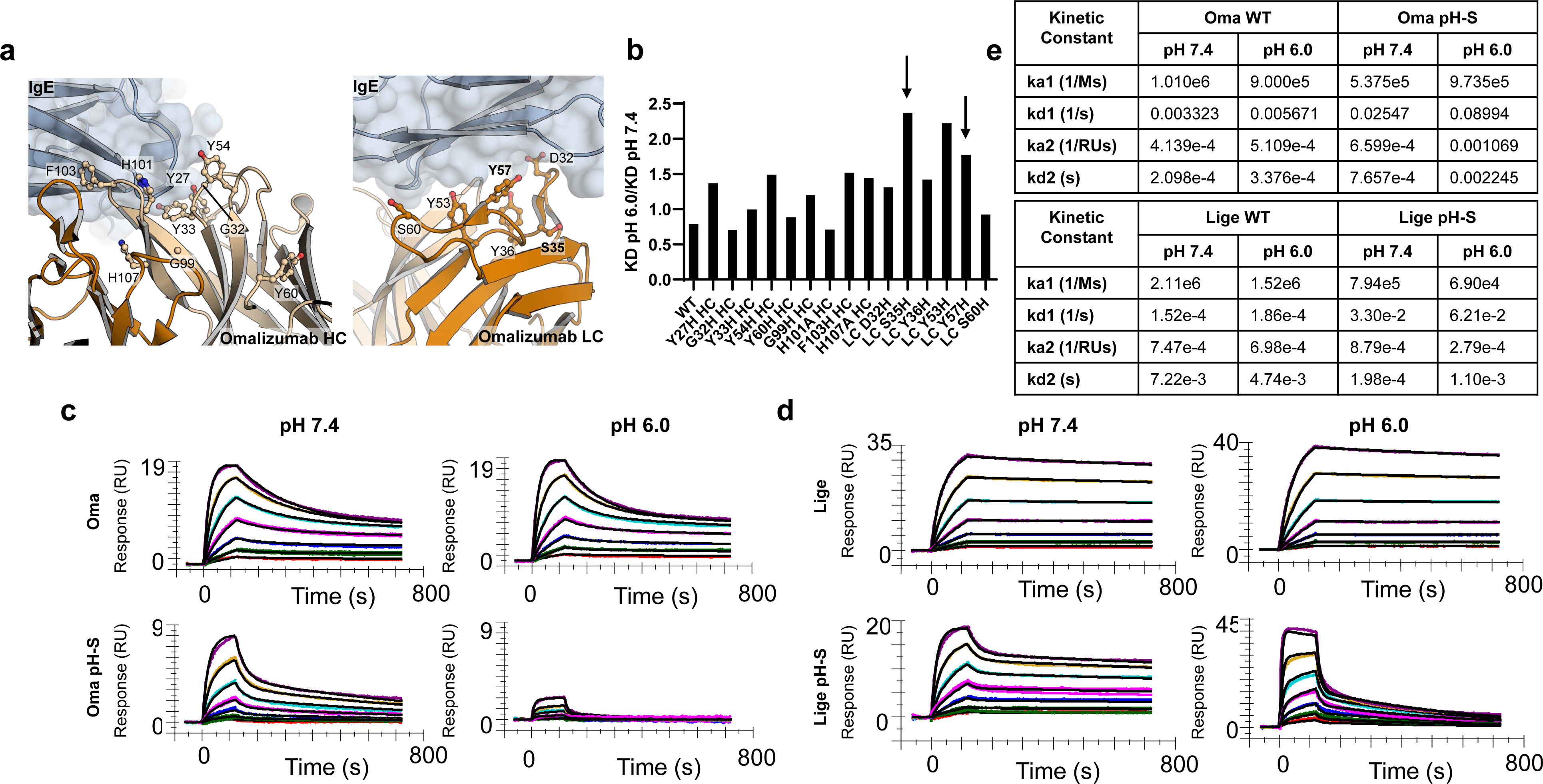
Design and characterization of pH-sensitive IgE binders. **(a)** Structures of omalizumab bound to IgE (PDB ID: 5HYS) highlighting heavy (left) and light (right) chain residues chosen for generation of single histidine point mutants. **(b)** BLI analysis of KD values at pH 6.0 versus 7.4; higher ratios indicate greater pH-sensitivity. Variants chosen for the generation of the Oma pH-S variant are highlighted with arrows. **(c-d)** SPR binding curves of omalizumab WT and pH-S (c) and ligelizumab WT and pH-S (d). **(e)** Table of binding constants obtained from SPR. Fits with a single site model were poor, so a bivalent analyte model was necessary to obtain meaningful parameters.

**Extended Data Fig 4.**
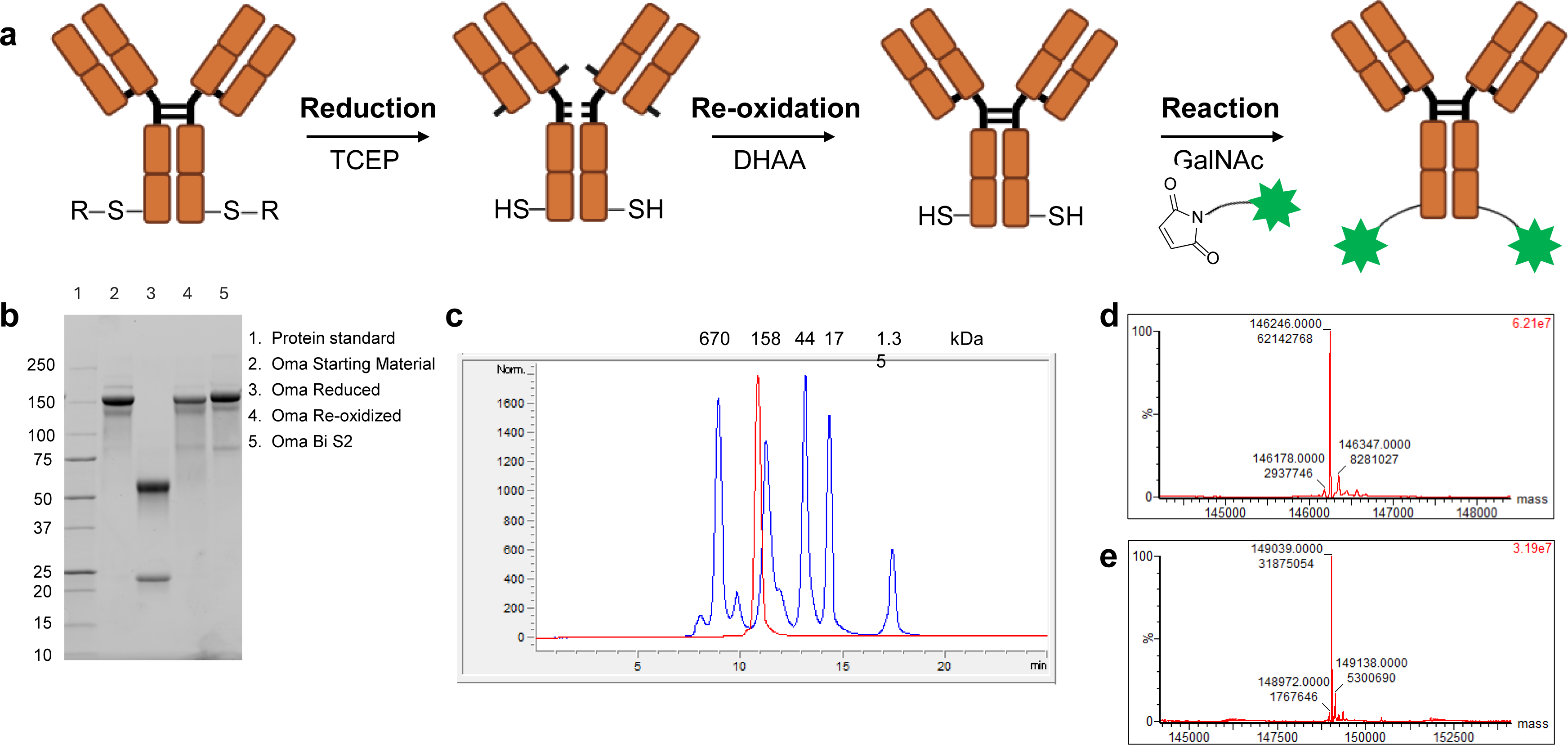
General conjugation strategy and characterization for GalNAc-LYTACs. (a) Antibodies engineered with the L443C cysteine mutation were labeled site-specifically with maleimide-GalNAcs. Antibodies were reduced with tris(2-carboxyethyl)phosphine (TCEP) to remove cysteine capping the L443C site, re-oxidized with dehydroascorbic acid (DHAA) to reform intramolecular disulfides, and the maleimide-GalNAcs were conjugated to free thiols on the L443C site. (b) SDS-PAGE was used to monitor Oma during the conjugation process. Precision Plus protein standard (Bio-Rad) was used to estimate mass. (c) HPLC size exclusion chromatography spectra of Oma Bi S2 demonstrates the conjugate is > 97 % monomer. Oma Bi S2 is shown in red, Gel Filtration Standard (Bio-Rad) is shown in blue. (d) Xevo-TOF-MS spectra of re-oxidized Oma. (e) Xevo-TOF-MS spectra of Oma Bi S2. The expected mass shift for the DAR 2 Oma Bi S2 LYTAC is 2791 Da. The observed mass shift between DAR 2 Oma Bi S2 and re-oxidized Oma was 2793 Da.

**Extended Data Fig 5.**
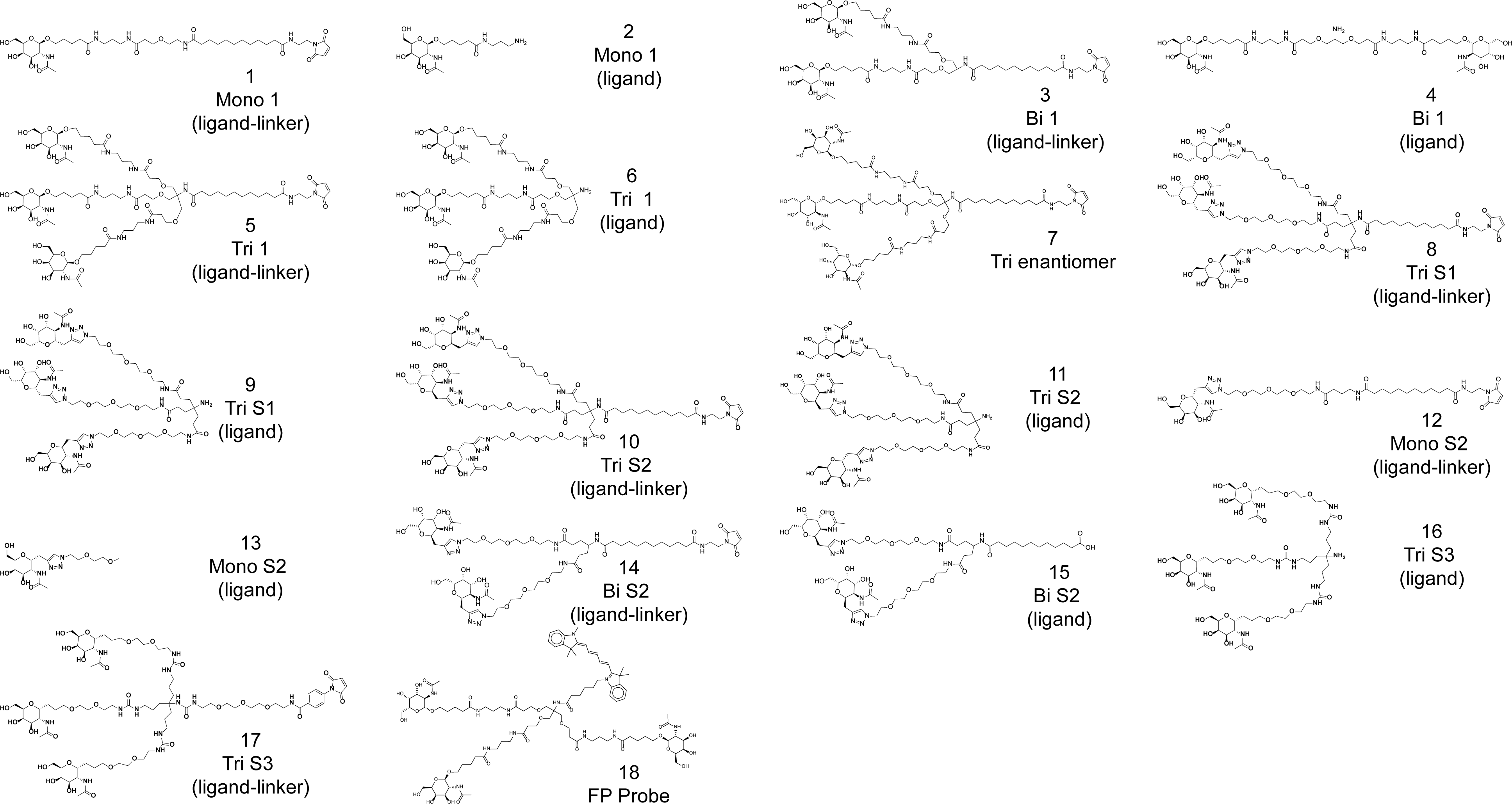
Structures of chemical ligands-linkers used for conjugation and ligand only molecules used for binding assessment in fluorescence polarization assays.

